# Agent-based Model for Microbial Populations Exposed to Radiation (AMMPER) simulates yeast growth for deep-space experiments

**DOI:** 10.1101/2023.10.29.564630

**Authors:** Amrita Singh, Sergio R. Santa Maria, Diana M. Gentry, Lauren C. Liddell, Matthew P. Lera, Jessica A. Lee

## Abstract

Space radiation poses a substantial health risk to humans traveling beyond Earth’s orbit to the Moon and Mars. As microbes come with us to space as model organisms for studying radiation effects, a computational model simulating those effects on microorganisms could enable us to better design and interpret those experiments. Here we present Agent-based Model for Microbial Populations Exposed to Radiation (AMMPER), which simulates the effects of protons, a major component of deep-space radiation, on budding yeast (*Saccharomyces cerevisiae*) growth. The model combines radiation track structure data from the RITRACKS package with novel algorithms for cell replication, motion, damage, and repair. We demonstrate that AMMPER qualitatively reproduces the effects of 150 MeV proton radiation on growth rate, but not lag time, of wild type and DNA repair mutant yeast strains. The variance in AMMPER’s results is consistent with the variance in experimental results, suggesting that AMMPER can recapitulate the stochasticity of empirical experiments. Finally, we used AMMPER to predict responses to deep space radiation that may be tested in future experiments. A user-friendly, open-source, extendable Python package for studying the relationship between single-particle radiation events and population-level responses, AMMPER can facilitate the basic research necessary to ensure safe and sustainable exploration of deep space.

## 1 Introduction

Space radiation poses a substantial threat not only to human explorers hoping to travel to deep space destinations such as the Moon and Mars^1^, but also to any organisms they bring with them. Much fundamental research remains to be done to understand and mitigate the biological effects of ionizing radiation on microorganisms, particularly in the deep space environment. Here we present AMMPER, a computational model for simulating the effects of deep space radiation on cells of the budding yeast *Saccharomyces cerevisiae*.

To date, most biological experiments in space have been carried out in Low Earth Orbit (LEO)^2^, on platforms such as the Space Shuttle and the International Space Station. However, the LEO radiation environment is fundamentally different from that of deep space (beyond LEO), the environment that will be experienced by crew in lunar or Mars missions, due to one key feature: Earth’s magnetic field^3–6^. The magnetic field protects spacecraft in LEO from exposure to a wide range of the high-energy particles that are present in the deep space environment^1^, including Galactic Cosmic Radiation (GCR) and Solar Particle Events (SPEs)^2^. Both GCR and SPEs are composed of highly energetic particles that cause DNA damage and oxidative stress, and exposure can compromise the viability and reproduction of cells, including microorganisms^1^. Organisms traveling to the Moon and Mars will experience both higher dose rates and a different spectrum of radiation types than on the ISS ^7–12^, making radiation one of the top 5 hazards to deep space astronauts recognized by NASA^13^.

Microbes are an integral part of human plans for deep space exploration for multiple reasons: they inevitably travel with us in the microbiomes of our built environments and our bodies, and they have the potential to play key roles in facilitating long-duration missions through functions such as biomining, life support, and production of food and pharmaceuticals^2,14–17^. Moreover, they are often favored as model organisms for space biology investigations due to their robustness to long periods of dormancy, ease of genetic manipulation, and relatively simple requirements for life support. A eukaryotic microbe, *S. cerevisiae* is a model commonly used to investigate human-relevant DNA damage and repair processes^18–20^.

Empirical data on the effects of deep space radiation on microorganisms are not extensive^2^, and are primarily limited to experiments on the survivability of cells in the dormant state^21–23^. However, exposure during dormancy omits the dynamic interaction between damage and repair characteristic of radiation-induced damage in actively growing and metabolizing cells. Biological CubeSats offer a solution by providing a relatively inexpensive platform in which microbes can be grown and monitored autonomously in liquid culture during spaceflight; these typically use optical measurements, where growth is measured by optical density and metabolic activity is measured by the reduction of the color-changing redox dye alamarBlue^24^. Biological cubesats have flown microorganisms both within LEO (e.g. GeneSat-1^25,26^, EcAmSat^27^, PharmaSat^28^) and beyond LEO (BioSentinel^18–20,29,30^), and the same technology will be used in NASA’s first experiment growing yeast on the Moon to investigate the effects of lunar gravity and radiation on biology (LEIA^31^).

Yet the very feature that makes CubeSat experiments more relevant makes their results more difficult to interpret: growth curve data from space-exposed microbes are an integration of the activity of thousands or millions of individual cells in suspension culture, of which some may have been hit by a charged particle, others by secondary radiation effects, and still others escaped unscathed. The temporally irregular and spatially heterogeneous nature of deep-space radiation remains an essential feature of the system^1,32^, but one that is undetectable in the final data. Reconciling these different scales will be essential both for predicting the performance of microbe-based space applications (e.g. fuel production in bioreactors) and for interpreting results from CubeSat experiments using microbes as model organisms. Agent-based modeling is one method that can allow researchers to explore how complex systems of individual parts such as radiation-exposed cells, each following their own rules, interact to generate population-level responses^33,34^.

A number of computational models exist for estimating biological effects of radiation; a brief overview is provided in the Supplementary Information. One model on which AMMPER builds is RITRACKS, which calculates dose due to radiation energy depositions in 3 dimensions^35^ (Fig. 1). Yet despite the number of existing tools, there are currently, to our knowledge, no models that analyze radiation effects from the perspective of a population of single-celled microbial organisms. Microbes are likely to show a very different response to deep-space radiation than do multicellular tissues, where chemical signaling can propagate a radiation-induced bystander effect, resulting in complex responses in cells never directly traversed by radiation particles^36–38^. Making effective use of microbes as human-relevant radiobiology model organisms therefore requires computational models that facilitate the interpretation of yeast radiation studies at the single-cell level. AMMPER (Agent-Based Model for Microbial Populations Exposed to Radiation) aims to fill this gap by simulating the effects of radiation on *S. cerevisiae* populations, using the ground experiments conducted for BioSentinel as a parametrization and validation case.

**Fig.1.**
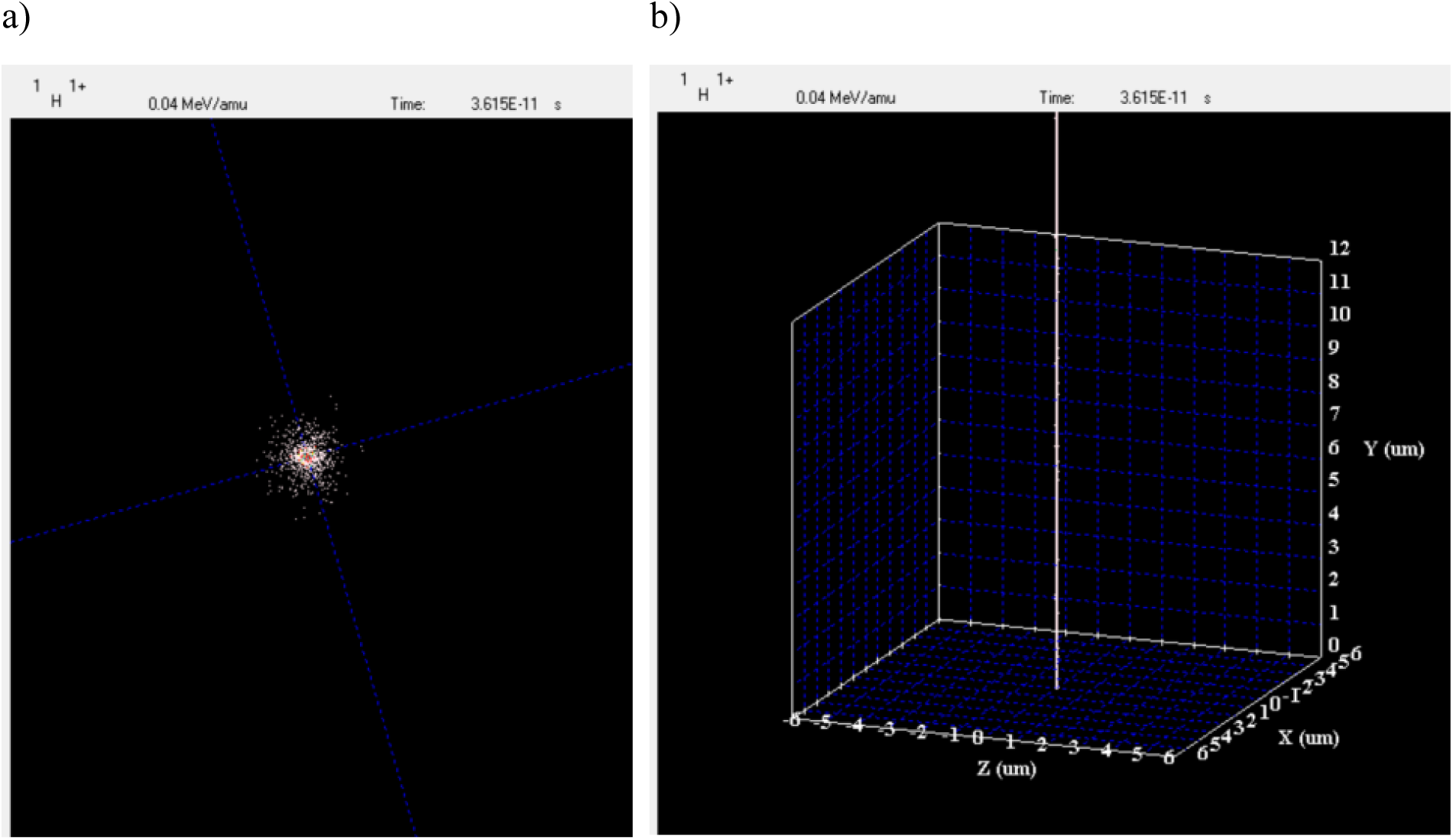
RITRACKS energy depositions^35^. Data generated from RITRACKS representing the energy depositions of a proton traversal, which passes through the point (0,0,0) and is oriented along the Y axis. (a) Cross-sectional view of proton traversal energy depositions. The density of electron energy depositions, indicated by the white dots, decreases radially further away from the proton traversal. (b) 3D view of energy depositions. Energy depositions (white dots) occur along linear proton traversal, and the radius of the track is small.

## 2 Results

### 2.1 AMMPER Overview

AMMPER integrates an existing model of energy deposition from particle radiation with novel algorithms for exponential population growth, DNA damage, DNA repair, and cell death in a single Python package (Fig. 2). By modeling the dynamics of a microbial population, AMMPER aims to generate testable predictions on the survivability of yeast in the deep space radiation environment.

**Figure 2.**
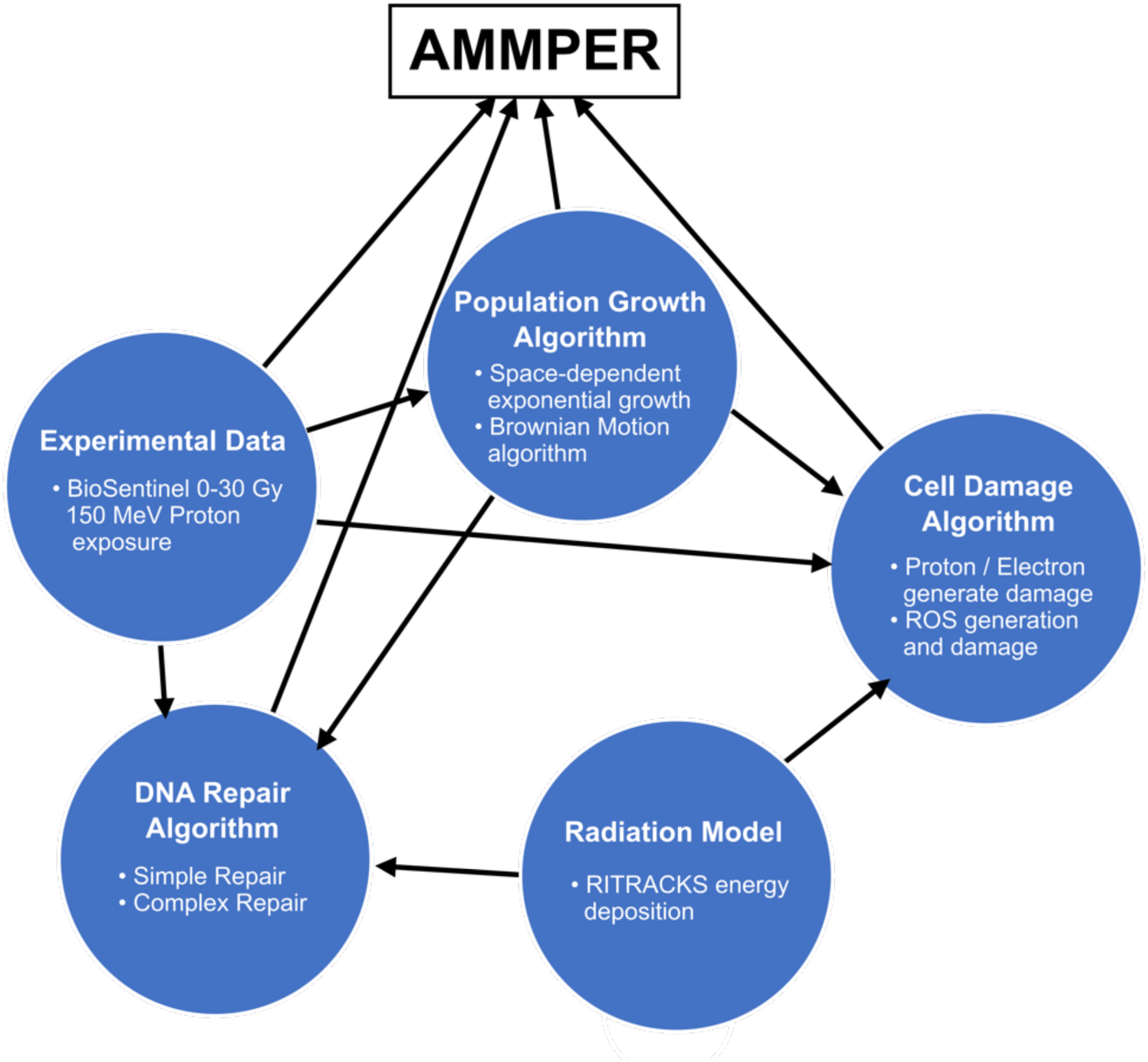
Block diagram of AMMPER. AMMPER integrates a model of radiation energy deposition with algorithms for population growth, cell damage, and DNA repair, and is parameterized with experimental data.

The AMMPER simulation space consists of a 64×64×64 µm cubic volume, analogous to an aqueous culture medium with non-limiting nutrient and pH buffering, in a typical laboratory microwell plate or a microfluidic culture card of the type used in BioSentinel^30^. The space initially contains a single model cell at the center that may then replicate to form a population (section 2.3). During population growth (for this study, we chose generation 2 as the exposure timepoint), a radiation environment is applied to the simulation space, consisting of an energy deposition map of both proton-based energy depositions and delta ray energy depositions (section 2.4). This energy map is used to calculate DNA damage in each of the cells, and to create Reactive Oxygen Species (ROS) molecules (section 2.4.3) that persist and continue to cause DNA damage and concentration-dependent apoptotic cell death in ensuing generations.

Based on parameters such as space availability and cell health, the model cells diffuse, replicate, and repair damage (sections 2.5-2.6).

### 2.2 Modeling Approach

AMMPER is intended to be a user-friendly tool to aid biological researchers with experimental design and data interpretation, and to assist in outreach and education on the effects of deep-space radiation. We therefore aimed to create the simplest model possible that could still recapitulate scientifically relevant features of experimental data, and designed it to be able to run on a standard laptop computer. In this initial release of AMMPER, the scientifically relevant features we targeted include the dose-dependent effects of proton radiation on population growth, and the variability observed among biological replicates of yeast growth experiments. Future versions of AMMPER may target different features to recapitulate.

AMMPER simulates a complex system consisting of physical, chemical, and biological processes. We began with information from subject matter experts and from the scientific literature to build an initial model capturing only what we hypothesized to be the salient features of these processes--those that would contribute to the features we wanted to recapitulate. The sources of the assumptions made for the cell growth, DNA damage, and DNA repair algorithms are described below (sections 2.3-2.6). We then carried out only two refinement steps: to determine the values of the doubling time parameter (section 2.3) and the ROS longevity parameter (section 2.3.3). Finally, we tested our hypothesis that this very simple model could generate similar effects to those observed in experimental data (section 2.8).

AMMPER is open-source software and publicly available through NASA GitHub^39^. The version presented here may be seen as an initial framework whose parameters and modules may be modified and improved upon in the future to accommodate other organisms and data types, and to simulate chemical and biological processes with better fidelity.

### 2.3 Model Cells

In AMMPER, each yeast cell is simulated as a sphere inscribed inside a 4×4×4 µm volume, resulting in a 33.51 µm^3^ cellular volume (Fig. 3) like that of a typical *S. cerevisiae* cell, with the nucleus sized to span 7% of the spherical volume^40^ (DNA is not modeled specifically). A cell agent has the following properties: UUID (Universally Unique Identifier), position of center, health status, generation at “birth,” generation at death, number of DNA Single Strand Breaks (SSBs), and number of DNA Double Strand Breaks (DSBs). The health status property has three options: 1 indicates a completely healthy (no DNA damage) cell, 2 indicates a damaged (nonzero amount of DSBs or SSBs) cell, and 3 indicates a dead cell^41^. Damaged cells are defined as cells that have halted replication until repair is complete, and dead cells are defined by their inability to repair damage or reproduce^42^.

**Figure 3.**
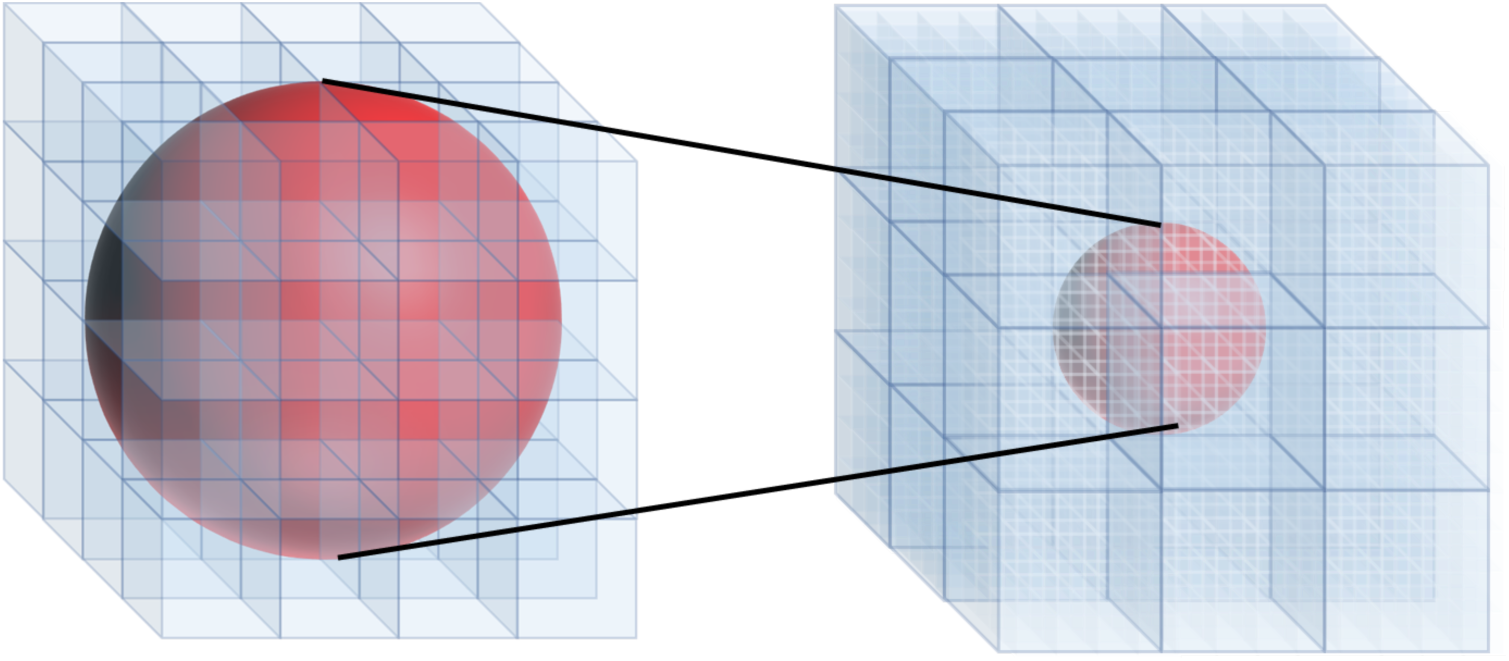
Model cell spatial definition. A cell is represented by the red sphere with a diameter of 4µm, inscribed inside a 4×4×4 µm cubic space (shown on the left, with each blue cube having dimensions of 1×1×1 µm). Each cell exists in a neighborhood (shown on the right, with each larger blue cube having dimensions of the 4×4×4 µm volume that holds a single cell), which consists of all 26 cell spaces bordering the cell’s cubic space.

AMMPER is implemented in generational time, with a single time step representing the amount of time it takes for a cell to replicate. We calculated the real-time value of one generation using empirical data from non-irradiated wild-type cells (see Methods, section 4.6); we found that the cell populations doubled in roughly 5 hours.

The cells exist in a three-dimensional neighborhood, defined by the 26 4×4×4 µm spaces surrounding it (Fig. 3). During replication, the cell object evaluates its neighborhood for empty spaces. Then, a random empty space is chosen for the cell to replicate into, at a rate of one new cell per generation. This form of replication results in exponential growth until saturation occurs, in which all 26 spaces around a cell are occupied and a cell can no longer replicate. To reduce the dependence of the growth curve on available space, a simplified Brownian motion algorithm is implemented that allows cells to spread out in space^43^. Like the replication algorithm, the cell object first evaluates its neighborhood for empty spaces. If no empty spaces are found, no movement occurs; otherwise, the cell moves into a random empty space. This algorithm increases the space between cells; the result is that saturation is delayed, and the population can maintain exponential growth for longer.

### 2.4 Radiation Environment

AMMPER has the capacity to simulate multiple GCR-analogous radiation environments: Deep Space Proton, NSRL GCRSim Proton^44^, and 150 MeV Proton. We chose to simulate 150 MeV proton exposure to correspond to the available empirical data. For Deep Space and GCRSim, this version of AMMPER simulates only the proton component for simplicity, because protons are the most abundant component of GCR: Hydrogen ions account for 87% of the total flux of GCR, with helium ions accounting for 12%, and other heavier elements (HZE) accounting for <1%^44^ (see Supplementary Background). While ions of higher Z may pose different health risks to multicellular organisms (an effect known as Relative Biological Effectiveness)^32^, very little data are available on the relative risks for single-celled organisms, so our initial focus is on ions with the highest chance of encountering cells. Future versions of AMMPER may include HZE as well as other low-LET radiation types, such as gamma radiation.

#### 2.4.1 Simulation Options

The Deep Space Proton model consists of the flux of the proton component of GCR, across the 90,000 µm^2^ cross sectional area of the 300×300×300 µm simulation volume, over 3 days in space beyond LEO^44^. The omnidirectional nature of space radiation is simulated by randomly generating traversal directions and positions over the simulation space^45^, and the dose rate of the Deep Space Proton model (4.49 mGy / 3 days) is intended to approximate the dose rate of protons present in the deep space environment. This is a unique feature of this computational model that is very difficult to reproduce in experiments. The NSRL GCRSim Proton model simulates the protons delivered during exposure to the NASA Space Radiation Laboratory (NSRL) Galactic Cosmic Ray Simulation (GCRSim)^44,46^, which provides exposure to a succession of energetic particles designed to simulate the major components of GCR. In AMMPER, GCRSim radiation is delivered during a short period of time and the traversals are oriented unidirectionally. Because one proton of each energy level is provided (for a total of 1940 MeV), the total dose delivered to the 64×64×64 µm simulation space is ∼1.19 Gy. The 150 MeV Proton model also contains unidirectional traversals but enables the user to vary the radiation dose delivered by 150 MeV protons to enable comparison to empirical data (Fig. 4). Detailed descriptions of the fluences for each simulation are included in Tables S1-S3 in the Supplementary Information.

**Figure 4.**
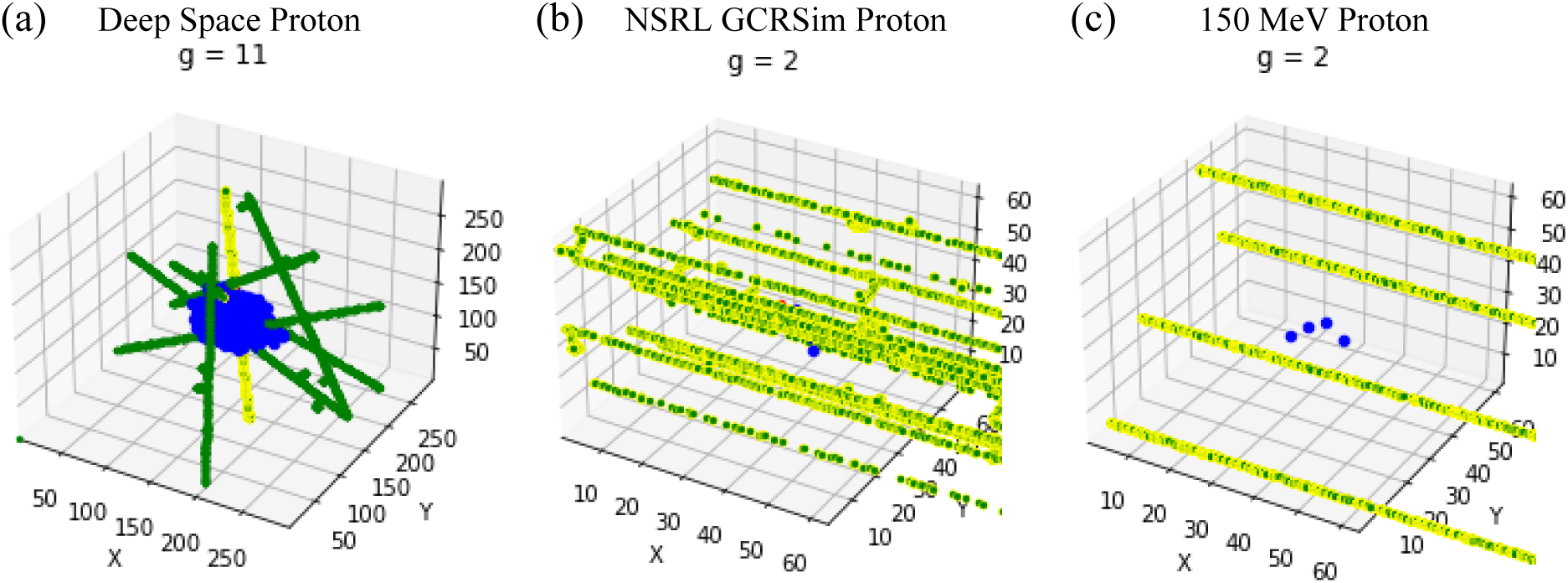
Radiation environments. Yellow dots represent energy depositions from proton and electrons, green dots represent ROS produced in the ionization of water molecules, and blue dots represent the cells in the simulation space. For all simulations, a cell is placed in the center of the cubic simulation space and undergoes exponential growth. (a) Deep Space Proton: The proton traversals are omnidirectional, and the timing of each proton is staggered to produce an environment that simulates the constant radiation exposure of deep space. Generation 11 shown here. (b) NSRL GCRSim Proton: All radiation is delivered at Generation 2, shown here. Proton traversals are parallel, and the total dose simulates the dose of a year of deep space radiation. (c) 150 MeV Proton, 10 Gy: All radiation is delivered at Generation 2, shown here. Total dose delivered is 10 Gy.

#### 2.4.2 Track Structure Calculations

To reduce simulation time, we precalculated the track structures resulting from individual proton traversals using RITRACKS^35^. We generated the energy deposition map for both ion and delta ray energy depositions at a resolution of 20 nm voxels. For each proton energy, we generated eight possible track options and unique energy deposition maps; the data for these are included with the AMMPER package. The absorbed dose from each track was calculated by summing all the energy deposition events associated with the track and dividing by the total mass of water in the simulation space, generating an estimate of the spatially-averaged absorbed dose from that track. To simulate a particular treatment, AMMPER code randomly selects tracks and combines the appropriate number of them to achieve the desired dose (Tables S1-S3). For the NSRL GCRSim Proton and the Deep Space Proton simulations, each track is also assigned a random position in space. Tracks for NSRL GCRSim are parallel to simulate the beamline source, whereas tracks for Deep Space Proton are randomly reoriented to simulate omnidirectional radiation.

As a confidence check, we compared energy depositions for proton energies between 0.5 to 1000 MeV in AMMPER to theoretical calculations for energy deposition as a function of radial distance from the particle track based on microdosimetry^47^ (Fig. 5a). We found that the microdosimetry calculation is consistent with AMMPER’s RITRACKS-generated data across the spectrum of 0.5 to 500 MeV protons. Additionally, the dose from delta ray energy depositions decreases as a function of radial distance from the core of the particle track (Fig. 5b), which is consistent with the findings of Wingate and Baum, who experimentally measured the radial distribution of dose from protons of energies between 1-3 MeV^48^.

**Figure 5.**
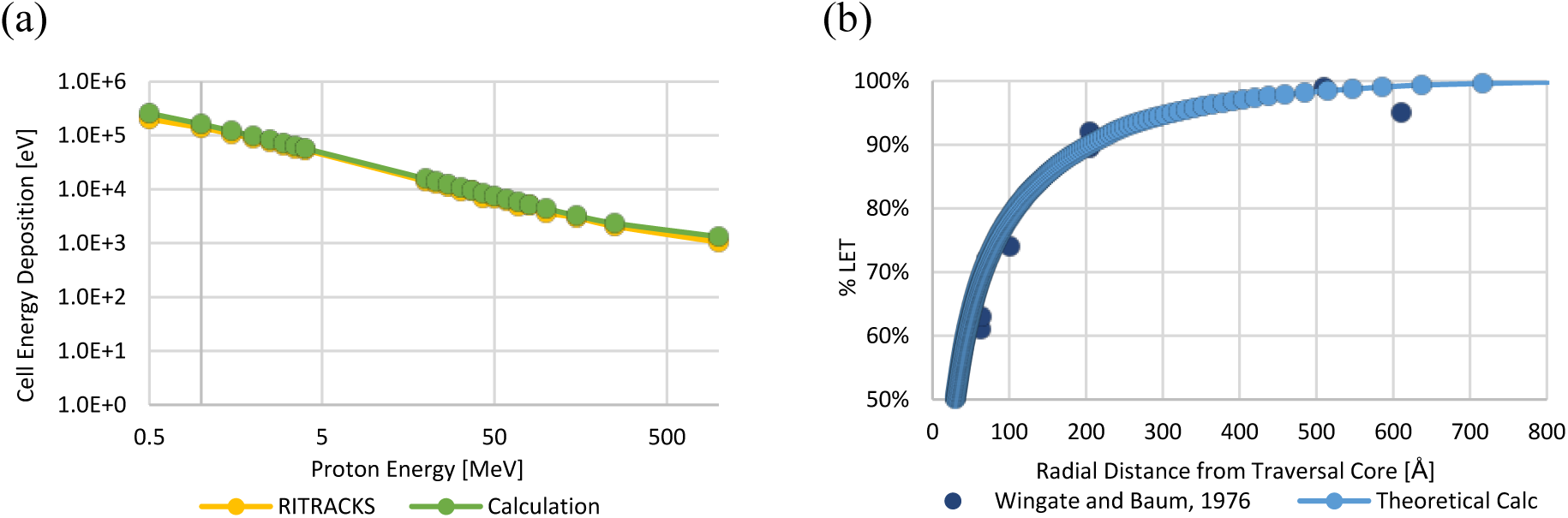
Distribution of RITRACKS ion and electron energy depositions. (a) Energy deposition in a single cell volume for radiation due to proton traversals of various energies. The RITRACKS data are depicted in yellow, and the microdosimetry-based calculations depicted in green. (b) Energy depositions by a 1 MeV proton. Light blue dots and line show energy depositions from delta rays, generated by RITRACKS, as a function of radial distance. Dark blue dots show Wingate and Baum^30^ experimental data on energy deposited by a 1 MeV Proton in tissue-equivalent gas as a function of radial distance. As distance from the proton track increases, the amount of energy deposited decreases.

#### 2.4.3 ROS Generation

We modeled the generation of ROS as a constant rate of OH**^·^** and H_2_O_2_ production, equivalent to the primary radioloytic yield of ROS from a 100 eV energy deposition in water (G_OH_ = 2.5, G_H2O2_ = 0.7 molecules/100 eV^49^). AMMPER’s native ROS generation algorithm calculates the amount of energy deposited in a 20 nm voxel, and, using the primary radiolytic yield of molecules / 100eV, calculates the number of molecules of OH**^·^** and H_2_O_2_ in that voxel (Fig. 6). For simplicity and speed of implementation in this initial version of AMMPER, we used the primary radiolytic yield of ROS in pure water as a simplified approximation for ROS generation rate. However, it is worth noting that the cell environment differs from pure water. Various elements (i.e. concentration of pH buffering, scavenging capacity of the cellular environment) affect these values^50^, and the model may be modified in future versions to take these elements into account, or, alternatively, to utilize the more detailed but computationally intensive chemical reaction calculations available in RITRACKS.

**Fig. 6.**
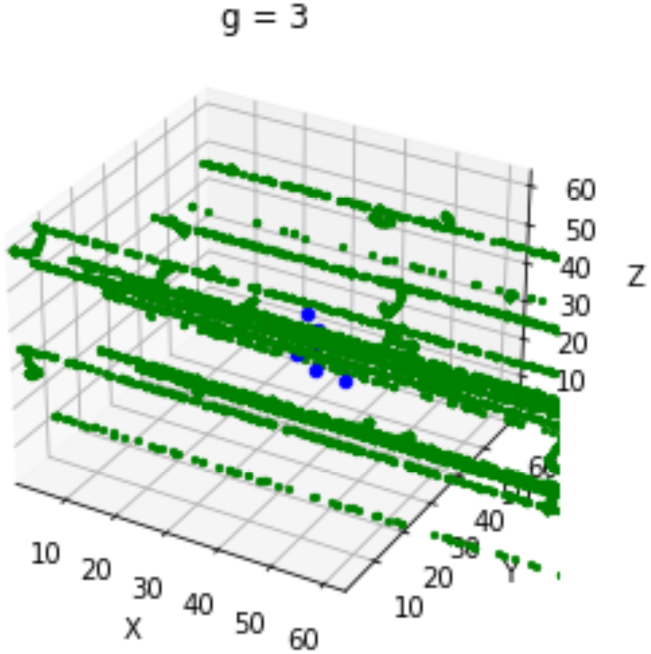
ROS species generation for the NSRL GCRSim Proton simulation in AMMPER. ROS specie,s are shown as green dots; each point corresponds to a single molecule of either OH^·^ or H_2_O_2_. ROS species are generated due to the ionization of water molecules by the energy depositions from the ions of the radiation tracks.

The longevity of ROS is complicated by ongoing chemical reactions and metabolic processes that result in the production of more ROS^51–54^. Because we were unable to find suitable experimental data on ROS lifetime inside yeast cells, we modeled ROS lifetime in AMMPER using a deterministic lifetime for each molecule, called the longevity parameter. For the initial implementation of the model, we chose a longevity parameter by assessing the effect of ROS longevity on yeast death in the model (Fig. 7).

**Fig. 7.**
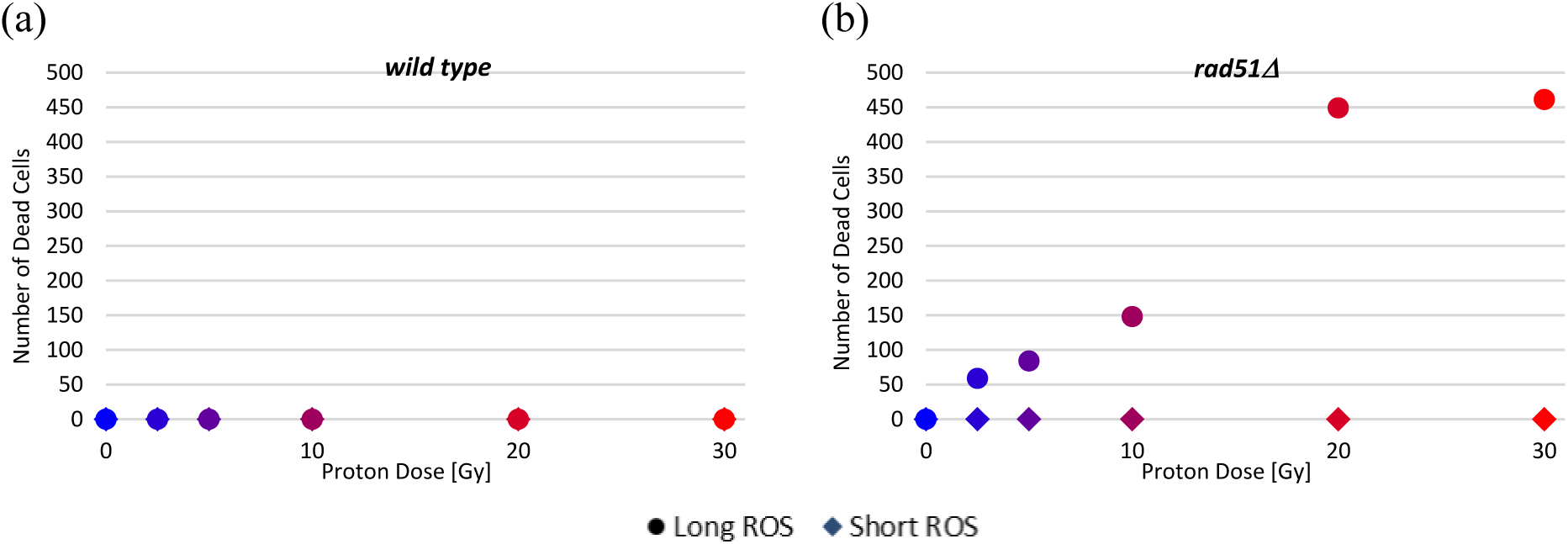
Comparison between cell survival for long and short ROS models. Total radiation dose in Gy is depicted along the x axis. The circles represent number of dead cells in the long ROS model, whereas the diamonds represent those in the short ROS model. Note that damaged cells are not included. Each symbol represents one model replicate. (a) Wild type simulated data. Circles and diamonds lie on top of each other; no cell death was observed in either ROS model. (b) rad51Δ simulated data.

We simulated two different yeast strains: wild type and *rad51*Δ, a DNA-repair mutant (see Methods). We compared the results of two ROS longevity parameters at the extreme ends of the range of potential values: “short” (1 generation, as short as possible) and “long” (>14 generations, or longer than remaining simulation time). In the simulated wild type population, both the long and short ROS models did not result in any cell death. In the simulated *rad51*Δ population, the long ROS model, but not the short ROS model, resulted in increased cell death as the radiation dose increased. The long ROS model is therefore necessary to generate cell damage or death. This long duration of ROS presence, which creates a “toxic environment” that persists in the space even after initial radiation exposure, may be interpreted as mimicking the complex chain of reactions that begin with the ionization of water molecules and result in long duration, non-targeted effects of ionizing radiation on cell populations^55^.

### 2.5 DNA and ROS-Mediated Damage

We implemented a simplified model of DNA damage and repair in AMMPER with the main goal of providing a quantitative structure to express a relationship between damage-causing elements (energy deposition and ROS contact) and effects on growth. A detailed consideration of the numerous forms of radiation-induced DNA damage and repair is beyond the scope of AMMPER (see Supplementary Background); for the model, we chose to focus on SSBs and DSBs, and to make a number of simplifying assumptions in order to keep the model tractable. We assumed that SSBs result primarily from electrons (delta rays), and DSBs only result from the actual proton traversal, as the energy required to generate a DSB is often higher than that contained in the delta rays^56^.

DNA is not explicitly modeled as a molecule in AMMPER; damage occurs due to energy deposition anywhere within the nucleus region. For each electron energy deposition at a nucleus, AMMPER generates an SSB, consistent with assumptions by Cucinotta *et al.* and Nikjoo *et al.* that SSB formation occurs when the energy deposition is above 17.5 eV^57,58^. AMMPER then calculates the radiation dose from proton energy depositions at the nucleus and determines the number of DSBs using an estimate of 35 DSBs/cell/Gy, assuming that there is a linear relationship between radiation dose and number of DSBs for *S. cerevisiae*^59,60^. Future versions of AMMPER may include more detailed models of DNA damage and repair (including types other than SSB or DSBs, like abasic sites, DNA cross-links, alkylation or oxidation of DNA bases) and/or simulations of different model organisms.

Under exposure to the simulated ROS environment, a cell may undergo additional damage. We assumed that each OH**^·^** molecule at the nucleus will cause an SSB. AMMPER also calculates the concentration of H_2_O_2_ in each cell to determine the probability of apoptosis, using the relationship between apoptosis and H_2_O_2_ concentration reported by Madeo *et al.*^61^.

If the number of DSBs or SSBs present in a cell is nonzero, the health status parameter increases to 2, indicating that the cell must halt the replication process until all damage is repaired. However, if apoptosis occurs, the health status parameter increases to 3, indicating that the cell is dead. Dead cells remain in the simulation space, but do not undergo the replication or repair algorithms^42^.

### 2.6 DNA Repair

In AMMPER, cells repair the different forms of DNA damage through two simulated mechanisms: “simple repair” and “complex repair,” which are distinguished by the time required for repair and probability of success^62–64^.

Simple repair is used on SSBs. In AMMPER, a cell repairs 3 SSBs per generation with a 100% probability of success. Complex repair is used on DSBs, and in AMMPER, a cell attempts repair on only 1 DSB per generation. Success is stochastic, with a 50% probability of success at each timestep, to account for the various probabilities of repair associated with the different DSB repair pathways, as reported by Lettier *et al.*^65^. When repair of a particular DSB is unsuccessful, the cell continues attempting to repair that DSB until successful before attempting repair on another DSB. Because genomes are not represented in this simulation, the cells do not have the capability to accumulate genome mutations. The values implemented in AMMPER for repair time and probability of success for the simple repair method are placeholders that we developed from the theory that base excision repair has a relatively high accuracy and low risk (see Supplementary Background), and were not refined or fit using experimental data. Additional research is required to determine the length of time it takes for a yeast cell to repair damage of various complexities.

### 2.7 AMMPER Outputs

After the simulation has finished running, AMMPER outputs data in comma-delimited a text file listing information from each timepoint on each cell present in the simulation space, and a folder of figures depicting the simulation environment, radiation traversals/timing, ROS generation, cell position, and cell health status at each generation of simulation time. These may be aggregated and processed by the user to generate data products as desired. For this study, growth curves were calculated from the number of dead, damaged, and healthy cells at each generation.

### 2.8 Empirical Data Comparison

As an initial evaluation of the ability of AMMPER to simulate the effects of radiation on microbial populations, we compared the data generated from AMMPER to those from a ground study done on yeast survival in response to ionizing radiation exposure that was carried out to support NASA’s BioSentinel mission. As described in section 2.2, the only components of the model that were refined using empirical data were the cell doubling time parameter; and the ROS longevity parameter. This comparison allowed us to test the hypothesis that the processes simulated in AMMPER are sufficient to recapitulate the dose-response relationship of two yeast strains to proton irradiation; and to generate variability among replicate simulations comparable to that observed among replicate experiments.

Two strains of *S. cerevisiae*, a wild type and a DNA repair mutant (*rad51*Δ; see Methods for details), were exposed to proton radiation at dose levels between 0 and 30 Gy. The cells were then allowed to grow, and optical density (OD_690_) measurements were taken for 83 hours. In AMMPER, *rad51*Δ cells were simplified to lack DNA repair capabilities entirely. Therefore, any DNA damage to the model *rad51*Δ cell type results in cell death, but in the absence of radiation their growth is identical to that of wild type. Our goal was to evaluate whether the model recapitulated two metrics from the empirical data: the relationship between growth rate and dose, and the variance among experimental replicates.

We found that there was no statistically significant dependence of growth rate on radiation dose in either strain in the empirical dataset, and AMMPER recapitulated this (Fig. 8; full growth curves in Fig. S2). In the *rad51*Δ strain, there did appear to be a decrease in growth rate as a function of the radiation dose at the lower dose levels (0-10 Gy), and an increase in growth rate at the higher dose levels (20-30 Gy), which were partially recapitulated in model results, but these trends were not statistically significant.

**Fig.8.**
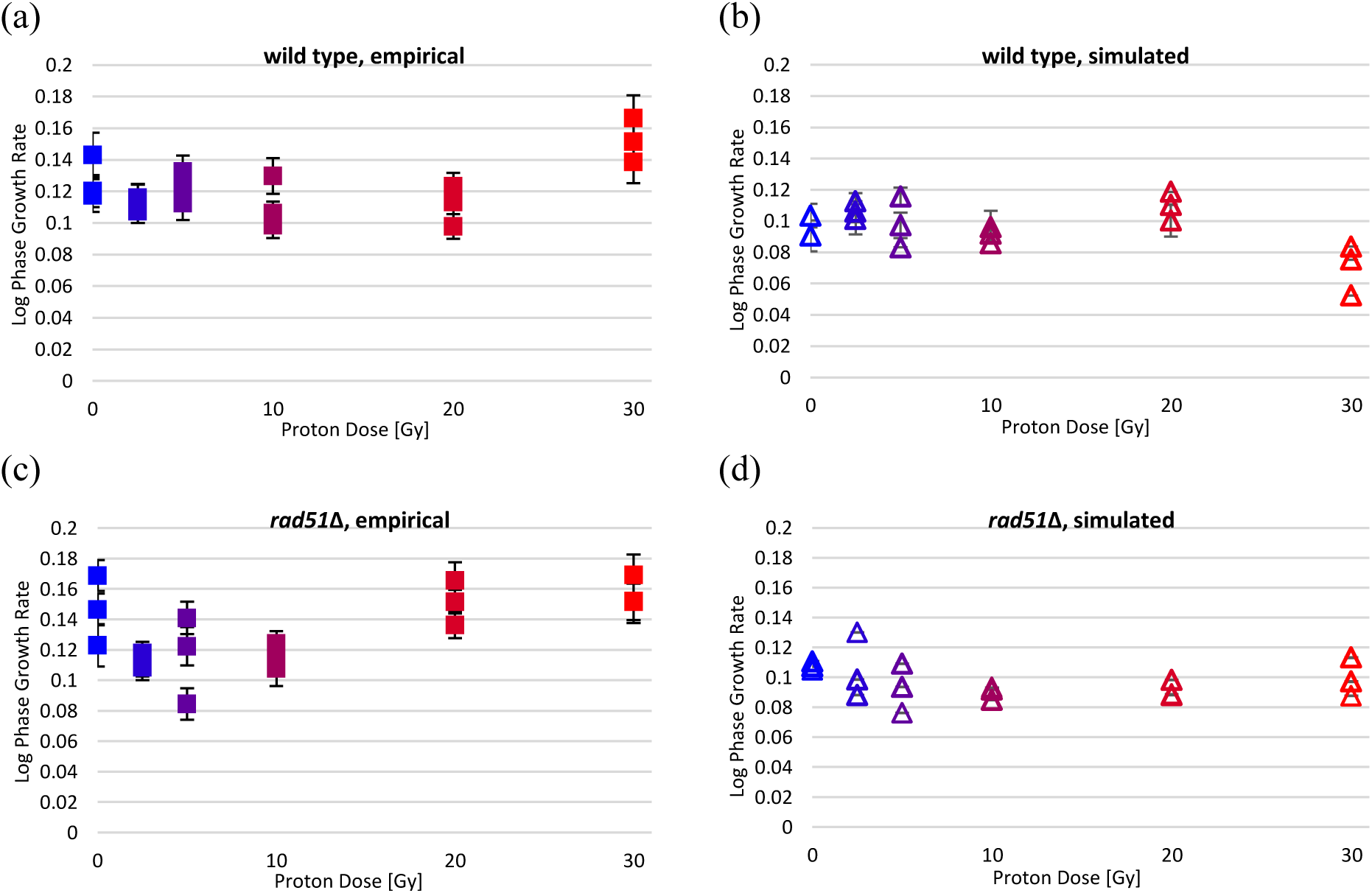
Comparison between empirical and simulated log phase growth rates. Radiation dose in Gy is depicted along the x axis. The open triangles represent the data from AMMPER, and the filled squares represent the empirical data. Each symbol represents one replicate (3 replicates/dose level for empirical and model data), and the error bars represent the linear regression fit error. Colors match the color codes in Fig. S2. (a) wild type empirical data; (b) wild type simulated data; (c) rad51Δ empirical data; (d) rad51Δ simulated data.

#### 2.8.1 Growth Rate

#### 2.8.2 Growth Rate Population Variance

Particle radiation is spatially heterogeneous, and its effects on a yeast population can vary widely depending on the overlap between particle tracks and the yeast cell locations. As an agent-based model, AMMPER can recapitulate some of this stochasticity. To measure this, we calculated the variance in the triplicate simulation and empirical growth rate data. We found that for both model and empirical data, variance among replicates was greater for the wild type than for the *rad51*Δ strain, but for both strains the variance among model replicates was of the same order of magnitude as in the empirical replicates. Variances were as follows: empirical wild type: 0.00157; simulated wild type: 0.00121; empirical *rad51*Δ: 0.000295; simulated *rad51*Δ: 0.000162.

#### 2.8.3 Lag Time

In addition to growth rate, lag time is a commonly measured parameter of microbial growth. Comparing the model and empirical results in lag time was not a primary focus of this study because 1) AMMPER does not explicitly model a physiological lag phase, so any apparent lag time would actually be a product of slow growth rate; and 2) the threshold for defining lag is different between simulation and empirical cases (simulation: number of cells; empirical: OD_690_), so the two lag times cannot be directly compared in absolute value. Nonetheless, we determined the relationship between lag time and radiation dose and compared between the empirical and simulated populations. The empirical wild type lag time did not exhibit a dependence on radiation dose; however, the empirical *rad51*Δ data displayed a statistically significant increase in lag time as the radiation dose increased, at levels below 30 Gy (slope: 0.5137; R^2^: 0.93, *p*: 0.0019; *p* < 0.05) (Fig. 9). Between 20 and 30 Gy, we observed a decrease in lag time, which may be related to the same complicating factors that also increased the growth rate in those samples. In contrast, the simulated data for both the wild type and *rad51*Δ strains did not exhibit a statistically significant dependence of lag time on radiation dose (Fig. 9).

**Fig.9.**
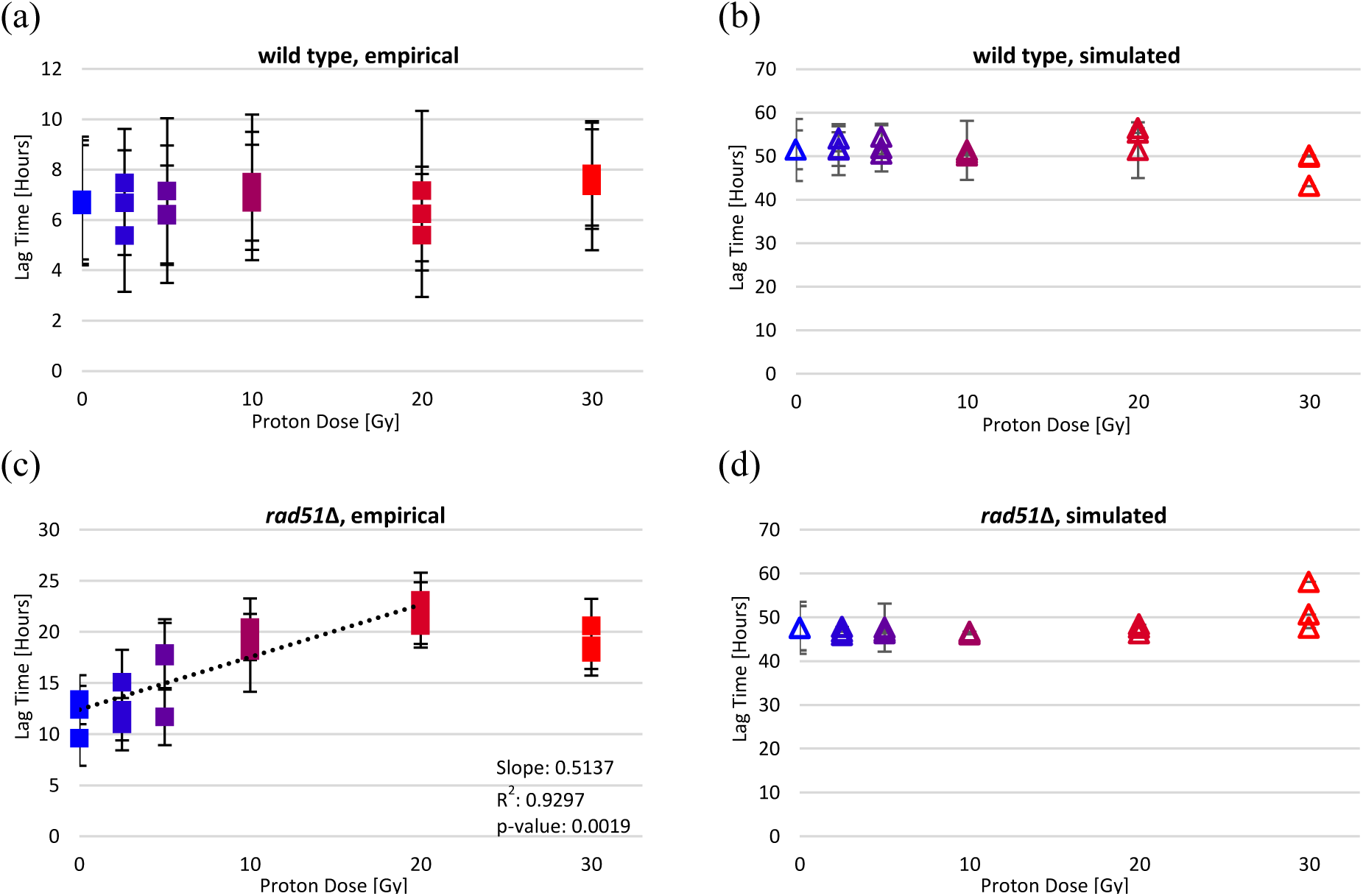
Comparison between empirical and simulated lag time. Radiation dose in Gy is depicted along the x axis. The open triangles represent the data from AMMPER, and the filled squares represent the empirical data. Each symbol represents one replicate (3 replicates per dose for both empirical and model data), and the error bars represent the error in the linear regression fit. Colors match the color codes in Fig. S2. (a) wild type empirical data; (b) wild type simulated data; (c) rad51Δ empirical data; (d) rad51Δ simulated data.

#### 2.8.4 Deep Space Proton/NSRL GCRSim Proton Simulations

One goal for the creation of AMMPER was to enable experimental design and hypothesis generation. To this end, we used the Deep Space Proton and NSRL GCRSim Proton models to predict results we might expect to see in future experiments (Fig. 10).

**Fig.10.**
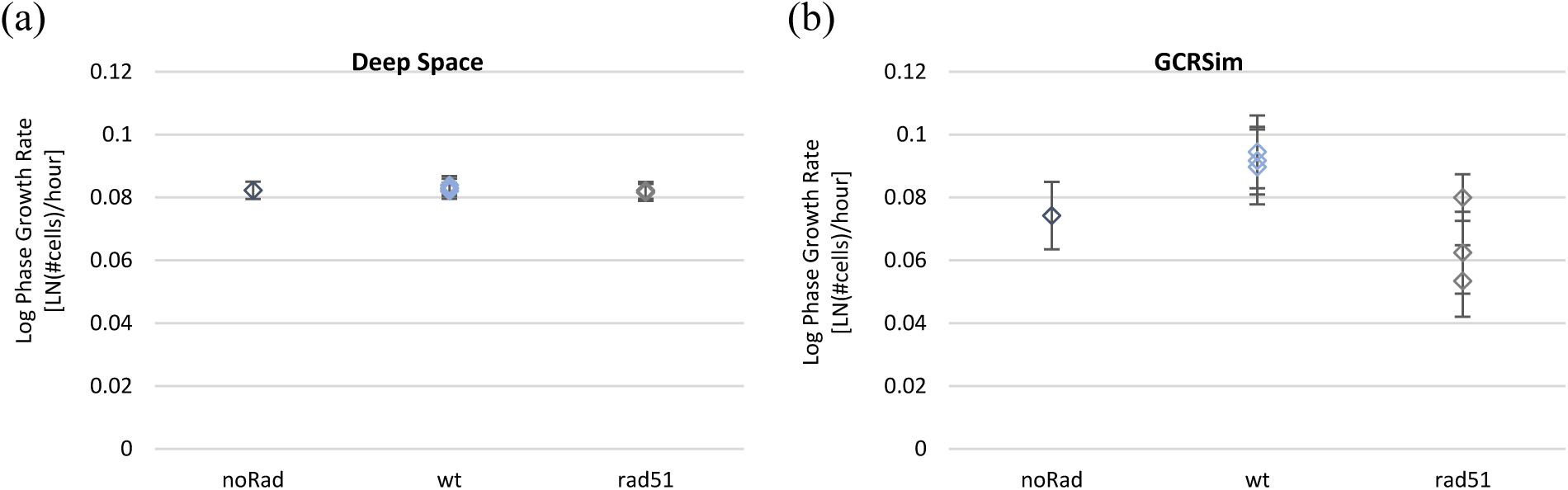
Log-phase growth rate for Deep Space Proton and NSRL GCRSim Proton simulations. Error bars represent the error in the linear regression fit. Dark blue, gray, and light blue represent the three replicates. noRad = control simulation without radiation, both strains (6 points). wt = wild type S. cerevisiae. rad51 = rad51Δ mutant. (a) Deep Space Proton simulation. There is minimal difference among the growth rates, regardless of modeled cell strain and radiation level. (b) NSRL GCRSim Proton simulation. Growth rate is higher in wild type cells exposed to radiation, and lower in rad51Δ cells exposed to radiation.

In the Deep Space simulation, cells experienced a fluence rate equivalent to what they would experience in deep space (see Table S1 for dose delivery), and the simulated three-dimensional space was 300×300×300 µm rather than 64×64×64 µm. This ability to simulate the very low dose rates of deep space is one of the major advantages of AMMPER, as it is currently not feasible to simulate deep-space radiation on Earth for experiments at realistic dose rates. The larger simulation size enabled us to observe more particle traversals for the same amount of simulation time, which is important given that the fluence rate in deep space is very low, although most traversals in the large simulation size do not interact directly with cells. Due to computational constraints, we were only able to simulate 3 days of growth for this project; future applications may include longer simulations. Growth rates during this time were unaffected by exposure to the deep space environment: we found no statistically significant differences among the control, wild type exposed to radiation, and *rad51*Δ cells exposed to radiation (ANOVA: df = 2, *F* = 2.41, *p* = 0.170) (Fig. 10). In contrast, the radiation delivered in the NSRL GCRSim simulation is equivalent to the dose and spectrum of particles that an astronaut would be exposed to over the course of a Mars mission, but delivered in the course of 1 hour^44^. We found that in this simulation, growth rates of irradiated *rad51*Δ cells were significantly lower than those of the irradiated wild type cells (ANOVA: df = 2, *F* = 8.87, *p* = 0.016; Tukey HSD for wt v. *rad51*Δ: q = 5.85 > q_crit_ = 4.34) (Fig. 10).

## 3 Discussion

With AMMPER, we have built a computational tool with the potential to model the activity of individual cells, to facilitate better prediction and interpretation of the population-level data generated by space radiation experiments with microorganisms. We have demonstrated that for a radiation environment composed of 150 MeV protons, AMMPER is effective at simulating the stochasticity exhibited in empirical data as well as the absence of radiation-induced decreases in growth rate at doses below 30 Gy.

As mentioned in the Introduction, AMMPER was built with simplicity as an initial priority, and it purposely includes the capacity for improvement and expansion. One difficulty we encountered in validating AMMPER was associated with the nature of the available empirical data. For this initial proof of concept, we focused on population growth of organisms exposed while in liquid culture as the most straightforward metric generated by the model. Future implementations of AMMPER may be modified to leverage data from a wider range of experiment types--for example, exposure during dormancy, heavy-ion particle damage on monolayers of immobilized cells^66,67^ or gamma-irradiated cells in continuous culture^68,69^ -- and from different metrics, such as the reduction of the metabolic dye alamarBlue, an assay commonly used as a more sensitive method for measuring non-lethal effects of spaceflight stress on microorganisms^19,27,70^. At the chronic low doses that are relevant to crewed deep-space travel, the effects of radiation on microbial cells may be primarily measurable as sub-lethal effects on metabolic activity, rather than as impaired growth or viability—making alamarBlue data especially important. Measuring and modeling these accurately will allow for a more accurate translation of results from microbial BioSentinel-like spaceflight experiments into estimates of long-term radiation risks to crew health.

In that vein, in the Deep Space Proton simulation we observed no effect of radiation dose on growth rate even for the sensitive *rad51*Δ strain. This is not unexpected, and is likely due to the low (chronic) dose rate. Prior spaceflight experiments with yeast have introduced very high dose rates in order to measure a response^71–73^. It has been estimated that an average yeast cell growing in deep space under 1 g/cm^2^ Al shielding could expect to see two months between proton traversals^1^, and actively growing cells have an advantage over dormant cells in that they can repair damage as it occurs. Future iterations of AMMPER will include other ions and neutral particles, as well as gamma rays, to enable more accurate representations of deep space as well as shielded and planetary surface environments. However, high-Z ions will encounter cells even less frequently than protons do. This observation is especially relevant in light of arguments being made by the radiobiology community on the importance of conducting more low-dose, chronic radiation experiments^46,74^. The BioSentinel mission includes several weeks to months of deep-space radiation exposure during dormancy prior to activation, increasing the probability of a radiation effect, but many anticipated applications of microbes in space biotechnology will not. Computational modeling tools such as AMMPER are therefore especially important in that they can allow researchers to explore the dynamic balance between damage rates and repair rates for different growth conditions, environmental conditions, and cell types.

Some observations from the empirical data lead us to believe that there may be missing components in the model whose inclusion would enable AMMPER to better represent biological processes. The fact that radiation elicited a dose-dependent increase in lag time in the *rad51*Δ strain not recapitulated by the model likely indicates that, unsurprisingly, radiation exposure perturbs cellular processes in ways not captured by a simple damage-repair-growth model. One unexpected result of the NSRL GCRSim Proton simulation was that radiation exposure appeared to increase growth rate in simulated wild type cells. This may be due to a complex relationship between space availability and growth rate that has still not been fully resolved in AMMPER. Additional improvements can be made to the DNA damage and repair models: DNA damage and subsequent repair pathways of *S. cerevisiae* have been heavily studied^75–78^, and chromosomal models of damage due to ionizing radiation already exist, such as those implemented in PARTRAC^79^. Implementing these higher-fidelity models of DNA damage could enable us to simulate single and clustered chromosomal aberrations, stalled replication forks^80^, and other complex features affecting the timing and probability of successful DNA repair.

AMMPER is a highly customizable simulation; each parameter mentioned in this article may be adjusted, each module of the model (yeast growth, DNA damage, repair, ROS dynamics) may be restructured or updated, and all underlying code is freely available to the public. Thus, AMMPER can be expanded to model diverse cell types and radiation environments to generate testable hypotheses about which components of the radiation response at the single cell level are the strongest determinants of the population-level effects. Indeed, we intend for the model published here to serve as a basic framework for more work to come. The ability of AMMPER to be modified to simulate a variety of dose rates enables the analysis of chronic exposure to radiation environments, which is not currently possible in traditional ground-testing experimental configurations. Furthermore, because AMMPER can represent stochastic results with a similar variance to biological data, it could be used to explore the proportion of variance due to different experimental components, and to assess what doses and dose rates would be necessary to obtain a statistically significant effect of radiation on growth rate. In later configurations, the capability to run Monte Carlo simulations could be implemented to enable analysis of the various outcomes in AMMPER.

In summary, AMMPER is a novel tool for mechanistic modeling of the biological effects of space radiation, using a flexible multi-parameter approach with simplified processes representing radiation energy deposition, ROS generation, DNA damage and repair, and cell reproduction. It tracks individual cell responses but enables comparison to population-level experimental data. We anticipate that future work with AMMPER will serve the space radiobiology community by supporting a greater mechanistic understanding of microbial radiation responses and aiding in experimental design.

## 4 Methods

### 4.1 AMMPER Development and Execution

AMMPER was written in Python 3.10.2, and utilizes packages included in the Python Standard Library, along with Numpy and Matplotlib. All model runs were executed on a PC laptop running Windows 10 with a 10th Generation Intel core i5-10310U processor and 16 GB of RAM. Simulations typically took approximately 30 minutes to run.

### 4.2 Microbial Culture Methods

In the ground study referenced here, wild type (YBS21-A) and *rad51*Δ (YBS29-1, *RAD51* gene deletion, necessary for homologous recombination-based DSB repair)^19^ diploid strains of *Saccharomyces cerevisiae* were exposed to 0 – 30 Gy 150 MeV protons at Loma Linda University Proton Treatment and Research Center (Loma Linda, CA). Strains were desiccated at 10,000 cells per well in 96-well Stripwell™ polystyrene microplates (Costar^®^; 9102) and kept in a dry state in cold storage (4°C) for 7 weeks. The cells were rehydrated with 100 microliters of Synthetic Complete (SC) growth medium^19^ directly before exposure to proton radiation, resulting in an initial density of 10^5^ cells/mL. Exposures were performed using a 20-cm diameter circular unmodified beam. The dose rates were 423.6, 844.8, 1,684.2, 3,367.2, and 5,051.6 cGy/minute for the 2.5, 5, 10, 20, and 30-Gy doses, respectively. Plate optical measurements (OD_690_) were initiated approximately 8 hours after exposure using a Molecular Devices VersaMax microplate reader, and measurements performed once per hour.

### 4.3 Model Initial Conditions

We designed the AMMPER simulation set-up to mimic the cell density of the experimental population at early log phase of an experiment, when OD_690_ = 0.3, corresponding roughly to a density of 3.85×10^6^ cells/mL. This density is equivalent to 2.5974×10^5^ µm^3^ per cell. Therefore, the AMMPER simulation was set to begin with a single cell at the center of a cubic 64×64×64 µm cubic space, which is equivalent to a volume of 2.62144 x10^5^ µm^3^/cell. A model simulation was conducted for each radiation dose three times. The radiation event occurred at generation 2, to approximate the experimental setup, described above, in which hydrated cells were irradiated in a single short treatment followed by several generations of growth. Simulated wild type and *rad51*Δ cells underwent log-phase growth under exposure to 0, 2.5, 5, 10, 20, and 30 Gy of 150 MeV proton radiation. Cell populations in a typical simulation typically reached 2-3 x 10^3^ cells per cubic space by the end of exponential phase (Fig. S2), equivalent to a density of between 7.63×10^9^ and 1.14×10^10^ cells/mL.

### 4.4 Track Randomization

To achieve a uniform distribution of the omnidirectional tracks, a spherical coordinate system was used. The theta and phi values (angular parameters for position in a spherical coordinate system) of the beginning and end of the particle track were generated randomly via a uniform distribution. The beginning and the end of the particle track were constrained to the surface of a sphere inscribed within the cubic simulation space. Then, a coordinate system transformation was carried out on the energy depositions of a pre-generated particle track to reorient it into the new position and direction.

### 4.5 Growth Rate and Lag Time Analysis

To evaluate AMMPER simulation outputs, we compared log-phase cell growth rates and lag times between simulated and empirical datasets. First, we manually selected the log-phase of the empirical data for both the wild type and *rad51*Δ *S. cerevisiae* populations. For the wild type samples, we determined that the log-phase occurred at 0.2-0.7 OD_690_ (Fig. S1a). For the *rad51*Δ samples, we determined that the log-phase occurred between 0.2-0.6 OD_690_ (Fig. S1b). We used the 0 Gy empirical data to calculate the doubling time of the population, which we then used to convert the simulation’s generational time to hours, facilitating comparison between the simulated and empirical data. For the simulation, growth rate was based on total cell count (rather than live cells only) for comparison with empirical OD data, which does not distinguish between live and dead cells. Lag time was defined as the time elapsed between the start of the experiment and when the growth curves reached OD=0.2 (for experiments), or between the start of the simulation and when the cell population reached 1000 cells (for AMMPER simulations). For the empirical data, “lag time” therefore includes contributions from both physiological lag (delay in growth due to cells’ transition out of stasis or recovery from damage) and differences in growth rate.

### 4.6 Statistical Analysis Methods

All statistical analysis was performed using Microsoft Excel. The growth rate for each simulated replicate was calculated using a least squares linear regression (LINEST, Microsoft Excel) for the number of cells at each timestep in hours. Each replicate was individually analyzed. The variance calculations for an assessment of the accuracy of the simulated dataset stochasticity was done using a Sample Variance function (VAR.S, Microsoft Excel). The comparison between the no radiation control group, *rad51*Δ, and wild type strains for the NSRL GCRSim Simulation and the Deep Space Proton Simulation was done using Tukey’s HSD in Microsoft Excel. The Anova: Single Factor analysis tool from the Excel Data Analysis ToolPak was used to determine whether the growth rates from the three groups differed significantly.

## 5 Code Availability

The code for AMMPER developed in this study is freely available on the NASA GitHub Repository (https://github.com/nasa/AMMPER)^39^.

## 6 Data Availability

The data generated by AMMPER in this study, and the empirical yeast growth data used in comparisons, are available in an online repository on Zenodo, DOI: 10.5281/zenodo.7379875.

## 8 Acknowledgments

We thank Robert Stewart, Sylvain Costes, and Kirtus Leyba for their assistance in refining the structure and model evaluation processes of AMMPER; Lynn Harrison, Marianne Sowa, and Egle Cekanaviciute for their contributions in how radiation exposure leads to radiobiological effects; Jared Broddrick and Tony Ricco for their discussions on how microbes function in the space environment; and Rocky An and Katie Blackwell for their support in troubleshooting various portions of AMMPER.

## 9 Author Contributions

A.S., J.A.L., and M.L. were responsible for model conceptualization, and S.S.M. provided input. A.S. carried out model development and coding, data analysis, and visualization. S.S.M., D.G., and L.L. generated the empirical data. A.S. was responsible for writing the initial draft of the paper; J.A.L., M.L., S.S.M., L.L, and D.G., reviewed and edited. J.A.L and M.L. were responsible for project management and mentorship.

