## Supplemental File 1 for "Agent-based Model for Microbial Populations Exposed to Radiation (AMMPER) simulates yeast growth for deep-space experiments"

### Supplementary background: Radiation models

There are several modeling methods that can be used to determine the risk of DNA damage due to ionizing radiation, including track structure analysis and microdosimetry[1]. Current computational models for the biological and chemical effects of radiation may be divided into various categories of focus: track structures, radiation chemistry, and intracellular chromosomal aberrations. One example from the track structures category is RITRACKS, which calculates a high-resolution, three-dimensional depiction of dose due to energy depositions from the ion itself (the core), as well as delta ray energy depositions extending radially outwards from the core particle track (the penumbra)[2] (Fig. 1). RITRACKS can also calculate radiation chemistry and chromosomal aberrations via DNA structure and damage simulations, but it does not include the capability to simulate the interactions of individual cells within a population[1]. The radiation chemistry category consists of models such as OREC[3], SPCS[4], and RADYLS[5], which use Monte Carlo modeling for the formation, diffusion, and reactions of chemical species created by the ionization of water resulting from particle tracks but do not have the capability to calculate the radiobiological effect of these molecules. The chromosomal aberrations category includes PARTRAC[6] and RADIFF, which create detailed simulations of radiation-induced DNA damage[6,7]. PARTRAC contains a suite of Monte Carlo simulations to model the radiation track structure and subsequent response both on a chemical and a chromosomal level[6]. RADIFF (RADical DIFFusion) was developed to calculate DNA double strand break induction on pBR322 plasmid DNA, focusing primarily on radicals created by electron traversals[7].

1. Nikjoo, H., O'Neill, P., Terrissol, M. & Goodhead, D. T. Quantitative modelling of DNA damage using Monte Carlo track structure method. *Radiat. Environ. Biophys.* **38**, 31–38 (1999).
2. Plante, I. & Wu, H. RITRACKS: A Software for Simulation of Stochastic Radiation Track Structure, Micro and Nanodosimetry, Radiation Chemistry and DNA Damage for Heavy Ions. in (2014).
3. Bolch, W. E. *et al. Monte Carlo simulation of indirect damage to biomolecules irradiated in aqueous solution: The radiolysis of glycylglycine*. ORNL/TM-10851, 6994843 http://www.osti.gov/servlets/purl/6994843/ (1988) doi:10.2172/6994843.
4. Turner, J. E. *et al.* Studies to link the basic radiation physics and chemistry of liquid water. *Int. J. Radiat. Appl. Instrum. Part C Radiat. Phys. Chem.* **32**, 503–510 (1988).
5. Hamm, R. N., Stabin, M. G. & Turner, J. E. Investigation of a Monte Carlo model for chemical reactions. *Radiat. Environ. Biophys.* **37**, 151–156 (1998).
6. Friedland, W., Dingfelder, M., Kundrát, P. & Jacob, P. Track structures, DNA targets and radiation effects in the biophysical Monte Carlo simulation code PARTRAC. *Mutat. Res. Mol. Mech. Mutagen.* **711**, 28–40 (2011).
7. Plante, I. A review of simulation codes and approaches for radiation chemistry. *Phys. Med. Biol.* **66**, 03TR02 (2021).

### Supplementary background: The biological effects of radiation

Because cells in the deep space environment are constantly exposed to galactic cosmic rays (GCR) (in contrast to solar particle events, SPEs), we focus here on the biological effects of exposure to this form of radiation. GCR consists of the nuclei of chemical elements and is thought to be created from explosive events outside of our solar system, such as a supernovae[1]. While elements from hydrogen to uranium can be present, elements of $Z > 26$ (elements heavier than iron) are much less abundant[2]. Hydrogen ions account for 87% of the total flux of GCR, with helium ions accounting for 12%, and other heavier elements (HZE) accounting for <1%[1]. During the solar minimum, with dose for reference field normalized to 500 mGy, it has been calculated that a cell nucleus with a cross section of 100 $\mu m^2$ would be traversed by hydrogen 126 times over the course of a year[1,2]. Therefore, 64.2% of the radiation dose results from protons[1]. Due to the abundance of protons in the GCR environment, we decided that AMMPER would focus on simulating the unshielded proton fluence of GCR in the deep space environment. The ionizing radiation of GCR can damage cells via both direct and indirect effects.

As high-energy protons pass through an aqueous space such as liquid microbial culture medium, the proton creates a unique track structure that consists of a core of energy depositions from the proton (resulting in the direct effects of this radiation) and a penumbra of energy depositions from electrons, or delta rays (that initiate the indirect effects of radiation)[3,4].

#### *Direct effects*

The direct effects of ionizing radiation consist of DNA damage, as reported by Cucinotta *et al.*[5], including DNA double strand breaks (DSBs: breaking of both DNA strands). When unrepaired or misrepaired, DSBs may lead to DNA mutations, carcinogenesis, and cell death[6]. Ionizing radiation may also induce DNA single strand breaks (SSBs: breaking of one DNA strand[7]. Nikjoo *et al.* claims that the clustering of multiple non-fatal SSBs, DSBs, base damage, and other damage complications lead to a higher probability of DNA misrepair and subsequent DNA mutations or cell death[7]; however, the effects of clustering of DNA damage is not modeled in AMMPER.

#### *Indirect effects*

The indirect effects of ionizing radiation occur both on an intracellular and extracellular level, resulting from the interaction between the cell and the products of water radiolysis[8]. The ionization of water due to energy depositions results in the generation of Reactive Oxygen Species (ROS), highly reactive molecules that include oxygen as a component. We focus here on those species created from an incomplete reduction of molecular oxygen, including hydrogen peroxide ($H_2O_2$), and the hydroxyl radical ($OH^{\cdot}$)[9]. The intracellular effects include the breakdown of biomolecules, resulting in a wide range of damage to various processes inside a cell[10], such as chemical modification to proteins, lipids, and nucleic acids[8]. On a cellular level, ROS generation leads to intracellular damage through oxidative stress, primarily resulting in necrosis[11]. Additionally, some ROS, such as $H_2O_2$, have been found to trigger apoptosis in *S. cerevisiae*[9,12–14]. Madeo *et al.*[12] tested for the induction of an apoptotic phenotype in exponentially growing yeast cells after exposure to various concentrations of $H_2O_2$, and found that at low concentrations, cell death occurs via a programmed apoptotic shutdown. Different concentrations of $H_2O_2$ result in different fractions of the cell population undergoing apoptotic death, with 0.3 mM $H_2O_2$ correlated with 20% of cells undergoing apoptotic cell death, 1 mM $H_2O_2$ with 40%, 3 mM $H_2O_2$ with 70%, and greater than 5 mM $H_2O_2$ correlated with a decrease in the fraction of cells undergoing apoptotic cell death due to an increase in the fraction undergoing necrosis[12].

The longevity of ROS in microbial culture medium is not well described in literature, as the presence of ROS may initiate several reactions that can lead to the production of additional reactive species[10,15,16]. $H_2O_2$ may also be generated through metabolic processes in cells, leading to further complications in determining the longevity of heightened concentrations of ROS[10]. The "bystander effect"[17], which Mothersill *et al.* defines as indirect and delayed responses of cells to low-dose radiation exposure, can result in mutations and apoptosis in unexposed cells[8], increasing the complexity of understanding and modeling the non-targeted effects of ionizing radiation.

1. Simonsen, L. C., Slaba, T. C., Guida, P. & Rusek, A. NASA's first ground-based Galactic Cosmic Ray Simulator: Enabling a new era in space radiobiology research. *PLoS Biol* **18**, e3000669 (2020).
2. Curtis, S. B. & Letaw, J. R. Galactic cosmic rays and cell-hit frequencies outside the magnetosphere. *Advances in Space Research* **9**, 293–298 (1989).
3. Cucinotta, F. A., Katz, R., Wilson, J. W. & Dubey, R. R. Radial dose distributions in the delta-ray theory of track structure. in *AIP Conference Proceedings* vol. 362 245–265 (AIP, 1996).
4. Cucinotta, F. A., Nikjoot, H. & Goodhead, D. T. The Effects of Delta Rays on the Numberof Particle-Track Traversals per Cell in Laboratory and Space Exposures. 5.
5. Cucinotta, F. A., Nikjoo, H., Wilson, J. W., Katz, R. & Goodhead, D. T. RADIAL DOSE MODEL OF SSB, DSB, DELETIONS AND COMPARISONS TO MONTECARLO TRACK STRUCTURE SIMULATIONS. 6 (1997).
6. Mladenova, V., Mladenov, E. & Iliakis, G. Novel Biological Approaches for Testing the Contributions of Single DSBs and DSB Clusters to the Biological Effects of High LET Radiation. *Front. Oncol.* **6**, (2016).
7. Nikjoo, H., O'Neill, P., Terrissol, M. & Goodhead, D. T. Quantitative modelling of DNA damage using Monte Carlo track structure method. *Radiation and Environmental Biophysics* **38**, 31–38 (1999).
8. Plante, I. Radiation chemistry and oxidative stress. 9 (2010).
9. Krumova, K. & Cosa, G. Chapter 1 Overview of Reactive Oxygen Species. 1–21 (2016) doi:10.1039/9781782622208-00001.
10. Jamieson, D. J. Oxidative stress responses of the yeast Saccharomyces cerevisiae. *Yeast* **14**, 1511–1527 (1998).
11. Sampson, T. R. *et al.* Rapid Killing of Acinetobacter baumannii by Polymyxins Is Mediated by a Hydroxyl Radical Death Pathway. *Antimicrob Agents Chemother* **56**, 5642–5649 (2012).
12. Madeo, F. *et al.* Oxygen Stress: A Regulator of Apoptosis in Yeast. *Journal of Cell Biology* **145**, 757–767 (1999).
13. Carmona-Gutierrez, D. *et al.* Apoptosis in yeast: triggers, pathways, subroutines. *Cell Death & Differentiation* **17**, 763–773 (2010).
14. Perrone, G. G., Tan, S.-X. & Dawes, I. W. Reactive oxygen species and yeast apoptosis. *Biochimica et Biophysica Acta (BBA) - Molecular Cell Research* **1783**, 1354–1368 (2008).

15. Attri, P. *et al.* Generation mechanism of hydroxyl radical species and its lifetime prediction during the plasma-initiated ultraviolet (UV) photolysis. *Sci Rep* **5**, 9332 (2015).
16. Thomas, J. K. Rates of reaction of the hydroxyl radical. *Trans. Faraday Soc.* **61**, 702 (1965).
17. Mothersill, C. & Seymour, C. Radiation-induced bystander effects: past history and future directions. *Radiat Res* **155**, 759–767 (2001).

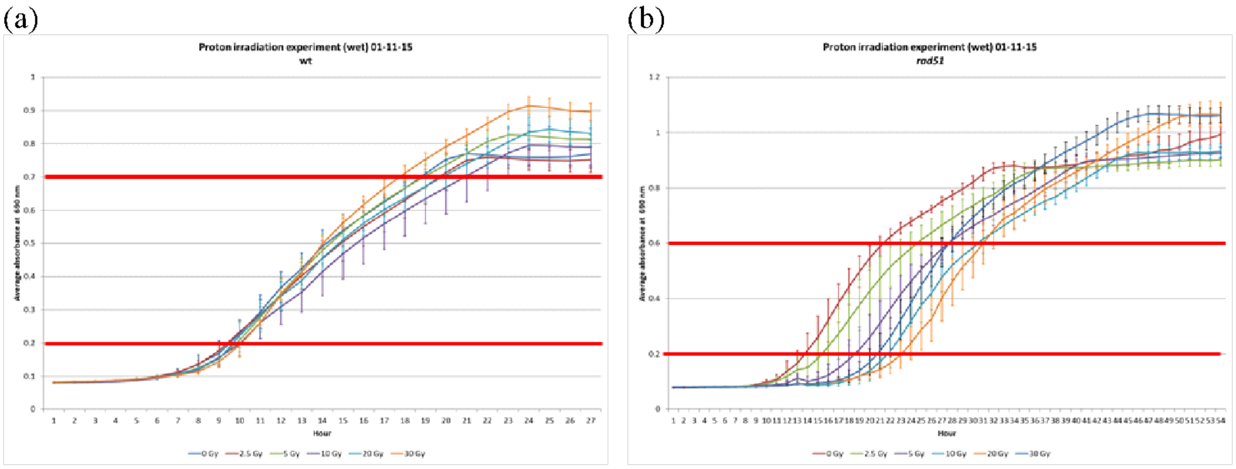


**Figure S1.** Growth curves of a) wild type and b) *rad51*Δ strains after radiation exposure (empirical data). Color denotes total dose (see legend). Data selected for growth rate calculations are those that lie between the horizontal red lines.

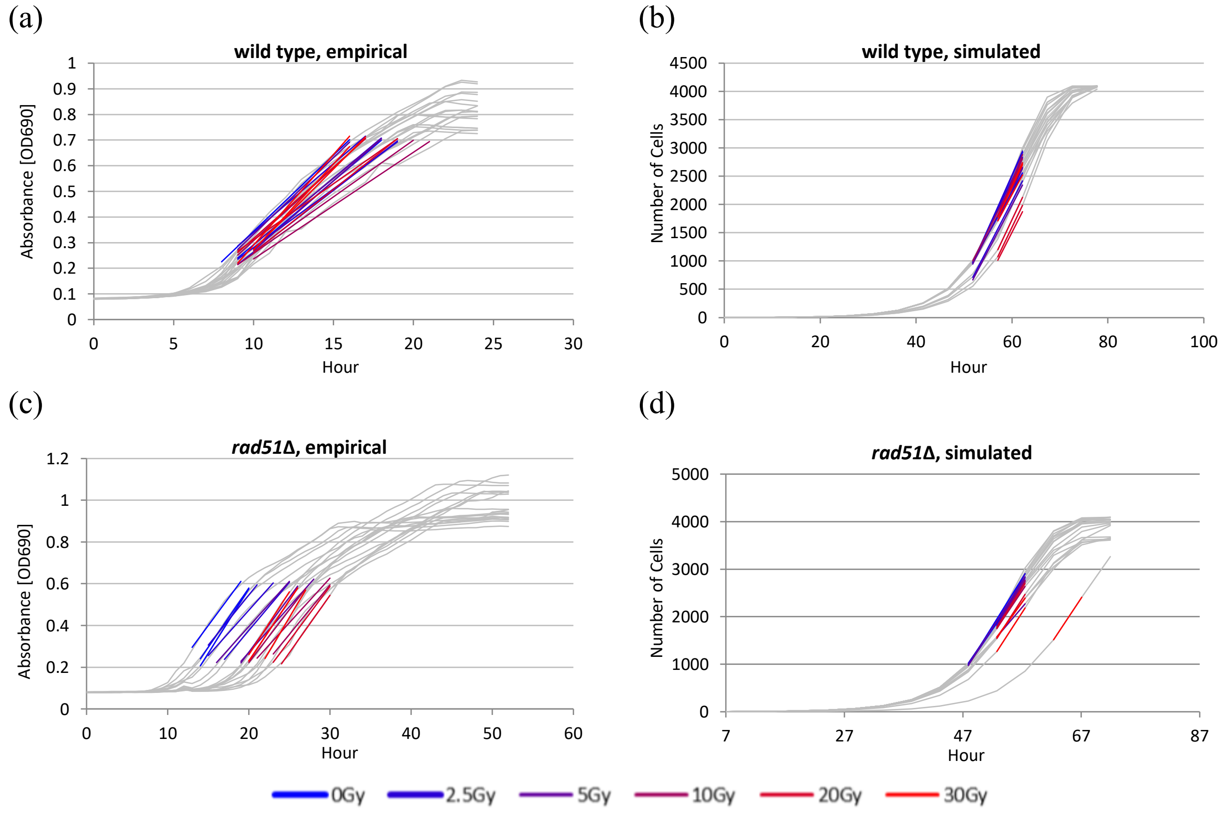


**Figure S2.** Comparison between empirical and simulated growth curves. Gray lines represent growth curves; colored sections represent the linear regression fit of the log phase portion of the growth curve. Radiation dose in Gy is depicted on a gradient from blue (0 Gy) to red (30 Gy). Each replicate is plotted individually. (a) wild type empirical data (b) wild type simulated data (c) *rad51*Δ empirical data (d) *rad51*Δ simulated data.

(a)

(c)

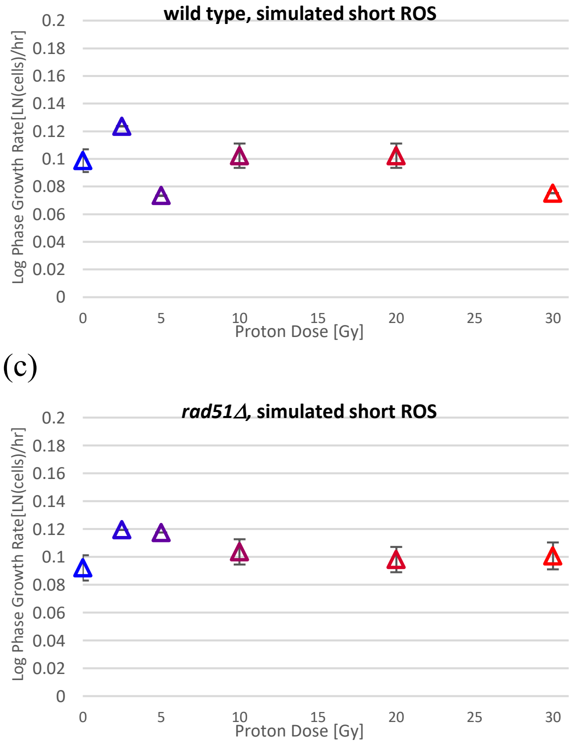


**Figure S3**. Short ROS simulation results. ROS longevity was adjusted for each molecule to disappear from the simulation space after 1 generation, instead than existing permanently, as it does in the long ROS simulations. Radiation dose in Gy is depicted on a gradient from blue (0 Gy) to red (30 Gy). (a) wild type empirical data (b) *rad51*Δ empirical data.

**Table S1: Radiation Delivery in of Deep Space Proton Simulation**

| Proton Energy [MeV] | # Traversals | Generation |
|---|---|---|
| 42.76 | 1 | 2 |
| 47.93 | 1 | 4 |
| 53.73 | 1 | 6 |
| 60.24 | 1 | 8 |
| 67.55 | 1 | 9 |
| 75.77 | 1 | 10 |
| 85.01 | 1 | 11 |
| 95.41 | 1 | 12 |
| 107.14 | 1 | 13 |
| 120.35 | 1 | 14 |

**Table S2: Radiation Delivery in of NSRL GCRSim Proton Simulation**

| Proton Energy [MeV] | # Traversals |
|---|---|
| 20 | 1 |
| 23 | 1 |
| 27 | 1 |
| 32 | 1 |
| 37 | 1 |
| 43 | 1 |
| 50 | 1 |
| 59 | 1 |
| 69 | 1 |
| 80 | 1 |
| 100 | 1 |
| 150 | 1 |
| 250 | 1 |
| 1000 | 1 |

**Table S3: Radiation Delivery in of 150 MeV Proton Simulation**

| **Radiation Dose [Gy]** | **# Traversals** |
|---|---|
| 2.5 | 1 |
| 5 | 2 |
| 10 | 4 |
| 20 | 8 |
| 30 | 12 |
